# Oropouche virus infects human placenta explants and trophoblast organoids

**DOI:** 10.1101/2024.11.16.623866

**Authors:** Christina Megli, Rebecca K. Zack, Jackson J. McGaughey, Ryan M. Hoehl, Taylor Snisky, Amy L. Hartman, Cynthia M. McMillen

## Abstract

Clinical and epidemiologic evidence from the recent outbreak of Oropouche virus (OROV) has demonstrated increased severity in clinical disease and adverse pregnancy outcomes including miscarriage, stillbirth, and neonatal demise. Serological evidence suggests vertical transmission of OROV may be responsible. OROV has not been studied in the context of pregnancy and has unknown ability to infect the relevant tissues of the maternal-fetal interface, which have anti-viral properties; therefore, the mechanisms of vertical transmission are unknown. We used polarized human trophoblast stem cell organoids and human placenta explants to demonstrate that OROV (BeAn19991) infects and replicates in human tissues of the maternal-fetal interface including chorionic villi and the microbial-resistant cell, syncytiotrophoblast. Viral replication is robust within the first 24 hours post infection, and tissues from earlier gestations may be more susceptible to infection. These data indicate tissues at the maternal-fetal interface are susceptible to OROV infection and may facilitate vertical transmission, leading to adverse pregnancy outcomes.

## Introduction

Oropouche orthobunyavirus (OROV) is an emerging virus transmitted by biting midges that was first identified in Trinidad and Tobago in 1955. Five years later, the virus was isolated from sloths in Brazil (1), and since then, it has caused over 80 outbreaks in South and Central America (2). Historically, OROV caused self-limiting febrile illnesses associated with joint stiffness, intense headaches, and pain; however, the recent 2024 OROV outbreak has revealed a previously unrecognized diversity of severe clinical outcomes including adult fatalities.

Possible cases of vertical transmission have been reported for the first time as well as adverse outcomes in newborns including microcephaly, stillbirth and neonatal deaths (3, 4). A recent case series has strongly suggested vertical transmission of OROV is associated with microcephaly in Brazil (4). OROV RNA and antigen were detected in neonatal samples and IgM was present in antepartum maternal samples and neonatal CSF and serum. These data suggest transplacental viral transfer may be a mechanism of vertical transmission.

Although only correlative data exists at this time, OROV may be the second virus within the Bunyaviridae family to be vertically transmitted in humans. Rift Valley fever virus (RVFV), a mosquito transmitted bunyavirus endemic in Africa and the Middle East, can be transmitted in utero. Two neonates, born from a mother recently infected with RVFV, had severe hepatitis upon delivery (5, 6). Both infants and their mothers had RVFV-specific IgM serum antibodies and at least one died from severe disease. Furthermore, women infected with RVFV are four-times more likely to have a miscarriage (7). Histopathological investigations have not been performed on placentas of suspected or confirmed cases of OROV or RVFV vertical transmission; consequently, the extent of infection and cellular targets of these viruses have not been identified clinically.

Obtaining clinical samples from patients infected with an emerging pathogen with potential to be vertically transmitted during pregnancy has multiple challenges, and thus we have developed laboratory models to aid in understanding placenta function and infection. The placenta is a multi-structured organ that functions both in nutrient transport and forms a protective barrier (8, 9). The fetal-derived placenta includes chorionic villi with direct contact with maternal blood and fetal membranes that are in contact with maternal decidua. Chorionic villi that are large, branch-like structures consisting of three types of trophoblast cells. At the tips of each villous are extravillous trophoblasts that invade into the decidualized uterus (i.e., decidua) and support implantation. Lining the villi is a large multinucleated cell, called syncytiotrophoblast (STB), that is formed after fusion of highly proliferative cytotrophoblasts (CTB). Physical and chemical (i.e., microRNA and interferon-λ) properties of STB provide resistance to microbial infection (10-13). Trophoblast organoids form when human trophoblasts are grown on beads and/or differentiated in culture. These organoids can properly polarize to have STB on the outer layer and CTB on the inner layer forming a villous-like structure that recapitulates the architecture of trophoblasts at the maternal-fetal interface (14). Three-dimensional trophoblast organoids provide a genetically tractable means to study viral replication at the maternal-fetal interface (15-18).

Surrounding the fetus are fetal membranes, which can be mechanically separated into chorionic and amniotic membranes. Vertical pathogens such as human cytomegalovirus, *Toxoplasma gondii* or Zika virus must traverse these layers to infect the fetus (19-22). Breaching these protective barriers and subsequent infection can lead to adverse pregnancy and/or neonatal outcomes (23-25). Human placenta explant cultures are valuable models to understand the local immune response and tropism of viruses, including Zika virus, HIV, and SARS-CoV-2 (26-28). Explant cultures have also played a key role in identifying cellular and tissue targets of the related bunyavirus RVFV at the maternal-fetal interface (29-31).

Recent reports of vertical transmission of OROV could be due to a change in virulence or a result of increased cases and heightened surveillance (32). Therefore, it is feasible that OROV may have undergone vertical transmission prior to the 2024 outbreak without recognition. To determine whether a historical strain of OROV infects human placentas and to identify which cell-types and tissues are targeted by the virus, we inoculated 3D human trophoblast organoids and human placenta explants with the prototype OROV strain BeAn19991. OROV replicated in both chorionic villi and amniotic membrane cultures, and earlier gestations were more permissive to infection.

## Results

To examine cellular tropism of OROV, we generated 3D trophoblast organoids from the trophoblast stem cell-derived cell line, CT-27, as previously described (33). These organoids were grown allowing for villous projections and development of the STB on the outer layer. After 5 days of growth organoids were inoculated with 1.0 × 10^5^ pfu of OROV BeAn19991 (34). Viral RNA was detected in the culture supernatant as early as 24 hours post-infection (hpi), which steadily increased by 48 hpi (**Figure 1A**). By 24 hpi immunofluorescence imaging showed organoid infection predominantly on the outer surface, whereas no virus was noted in mock infected controls (**Figure 1B & Supplemental Figure 1**). These findings highlight the susceptibility of key cells at the maternal-fetal interface to OROV infection.

**Figure 1:**
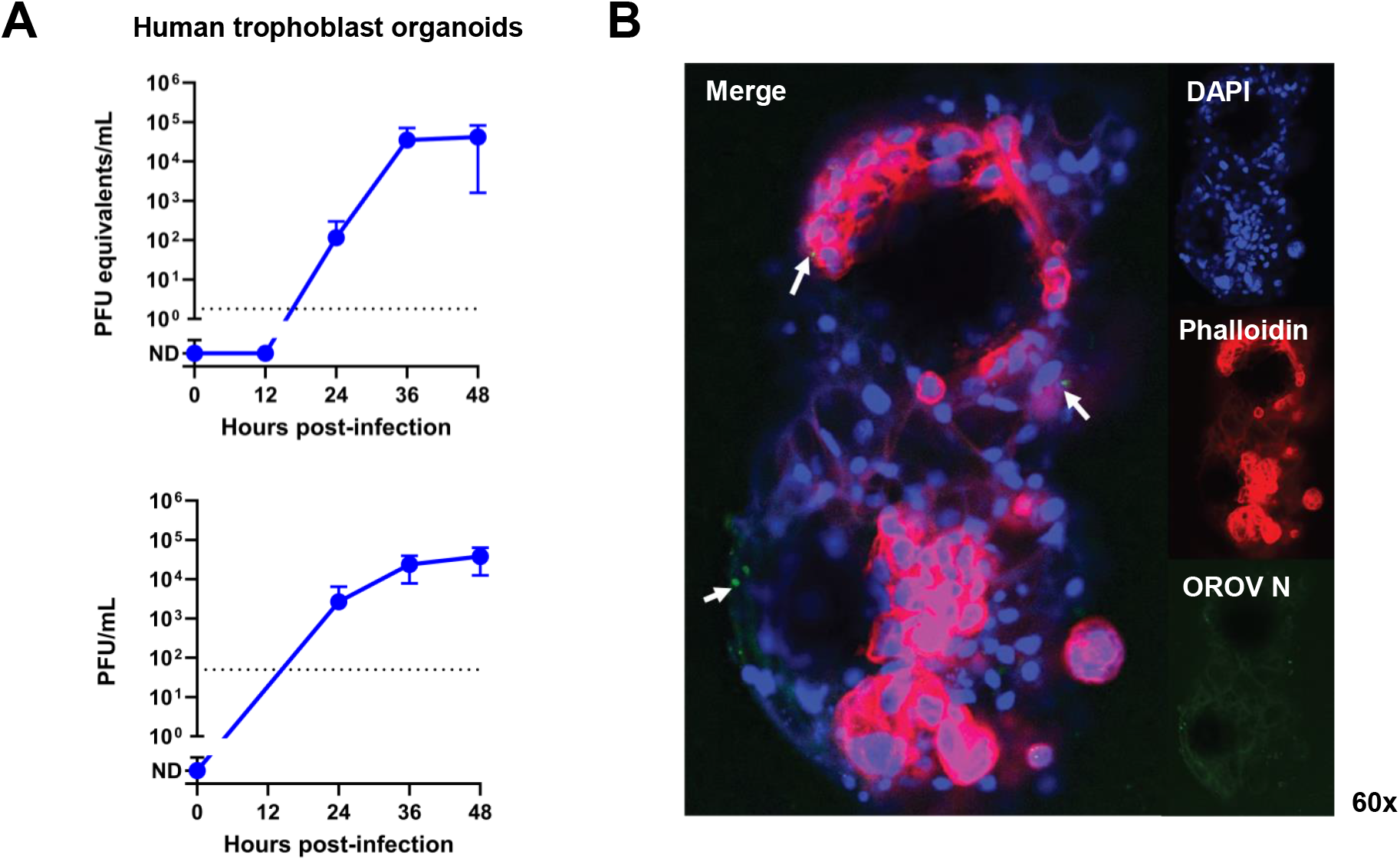
Oropouche virus replicates in trophoblast organoids. Viral growth curve of 3D trophoblast organoid cultures derived from differentiated CT-27 cells. Organoids were inoculated with 1.0 × 10^5^ pfu OROV BeAn19991, virus was removed and washed prior to addition of D2 culture medium, then culture supernatant was collected at 0, 12, 24, 36, 48 hours post-infection. **(A)** qRT-PCR was performed to quantitate virus production over time. Dashed line = limit of detection (LOD). Data represents one independent experiment with four biological replicates. (**B)** Immunofluorescence images of OROV infected organoid. blue = DAPI, red = phalloidin, green = OROV nucleoprotein (N). Image was taken at 60x magnification.

To evaluate the tissue and cellular tropism of OROV at the maternal-fetal interface, human placentas collected at 32-, 35-, 36-, and 39-weeks gestation were cultured as explants. Each placenta was separated into villous, chorionic membrane, amniotic membrane, fetal membrane (containing chorionic and amniotic membranes) and decidua, then dissected into 5 × 5 mm sections. Tissue dissections were inoculated with 1.0 × 10^5^ pfu of OROV BeAn19991 for 72 hours. The fetal membranes, including the chorionic and amniotic membranes, and decidua were not available for 32-week gestation placentas. A viral growth curve generated from culture supernatant showed sustained viral replication in villous cultures from placentas of fraternal twins at 32 weeks gestation and a 35-week gestation placenta (**Figure 2A**). By 72 hpi, 3-fold and 65-fold or 34-fold more infectious virus was detected than 0 hpi, respectively. Significant replication was not observed in the villous cultures from 36- or 39-week gestation samples. To validate infection of chorionic villous explants, we performed immunofluorescence (**Figure 2B**) and immunohistochemistry (**Figure 2C**) using antibodies to OROV nucleoprotein and dsRNA. Viral staining was noted in both the superficial STB layer as well as deeper tissue.

**Figure 2:**
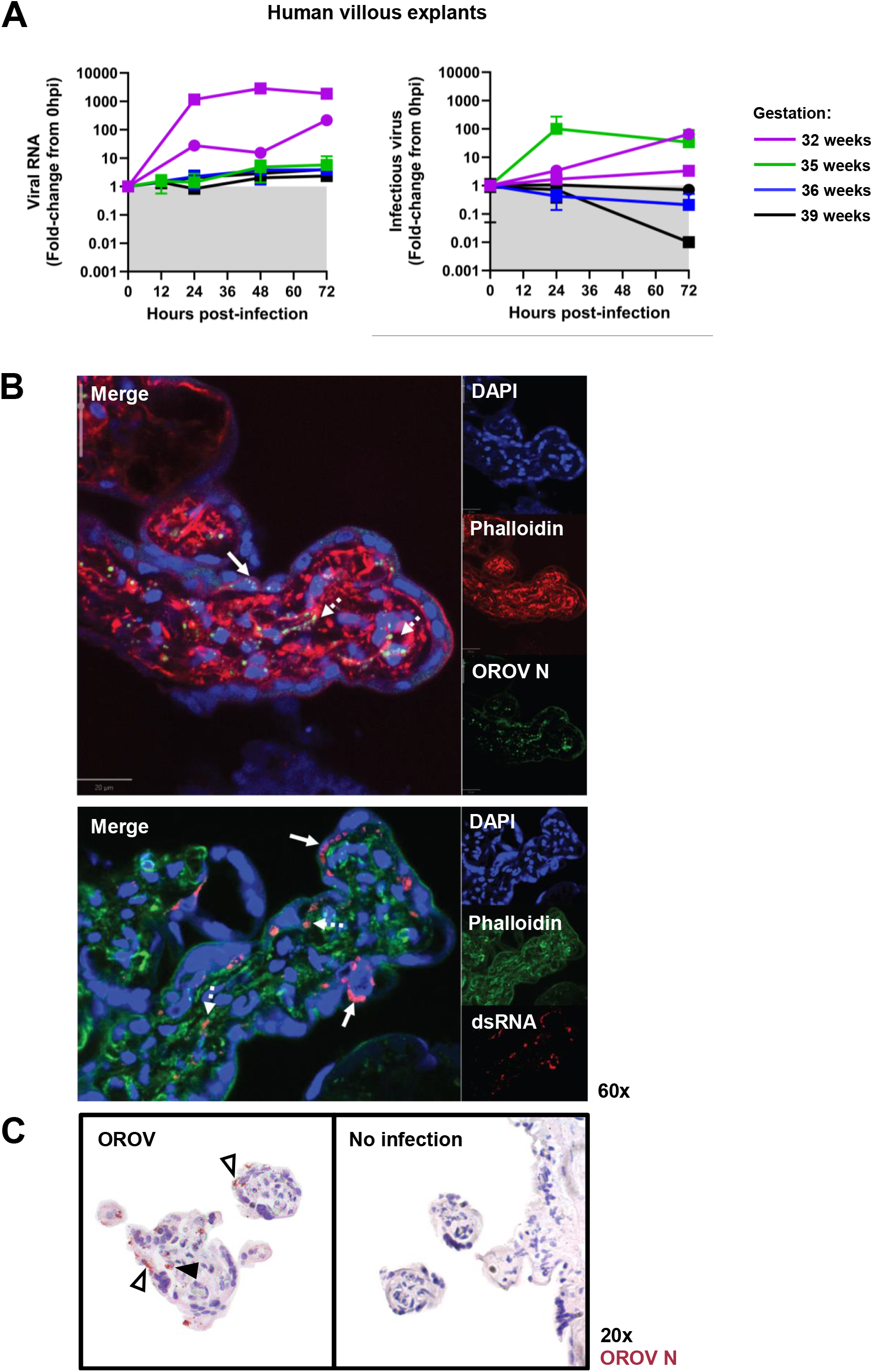
Oropouche virus infects and replicates in human villous explants and syncytiotrophoblast. Placentas were donated at 32- (n=2), 35- (n=1), 36- (n=1), and 39-week (n=2) gestation. Cultures of dissected decidua, villous, chorionic membrane, amniotic membrane, and fetal membrane were inoculated with 1 × 10^5^ pfu of OROV BeAn19991 for 1 hour (n=1-3 per donor). Virus was removed and washed prior to the addition of culture media. Culture supernatant was collected at 0, 12, 24, 48, and 72 hpi. **(A)** Q-RT-PCR and viral plaque assay was performed on culture supernatant to quantify viral RNA (left) and infectious virus (right) production over time. Dashed line = LOD. **(B)** Immunofluorescence of OROV infected chorionic villi (36-week gestation) demonstrates staining both superficially in the syncytiotrophoblast layer (solid arrows) and deeper in the villous tree (dashed arrows). Images were taken at 60x magnification **(C)** Immunohistochemistry of OROV infected chorionic villi demonstrate OROV in the superficial syncytiotrophoblast layer (open arrowheads) and deep in the center of the villous tree (closed arrowhead). Blue = DAPI, red = phalloidin (top) or dsRNA (bottom), green = OROV nucleoprotein (N; top) or phalloidin (bottom). Images were taken at 20x magnification.

OROV also replicated in fetal membranes (**Figure 3**). Amniotic membrane cultures from 35- and 36-week placentas had productive OROV replication over 72 hours (1100- and 61-fold increase, respectively). 36-week fetal membrane had approximately 50-fold more virus by 72 hpi.

**Figure 3:**
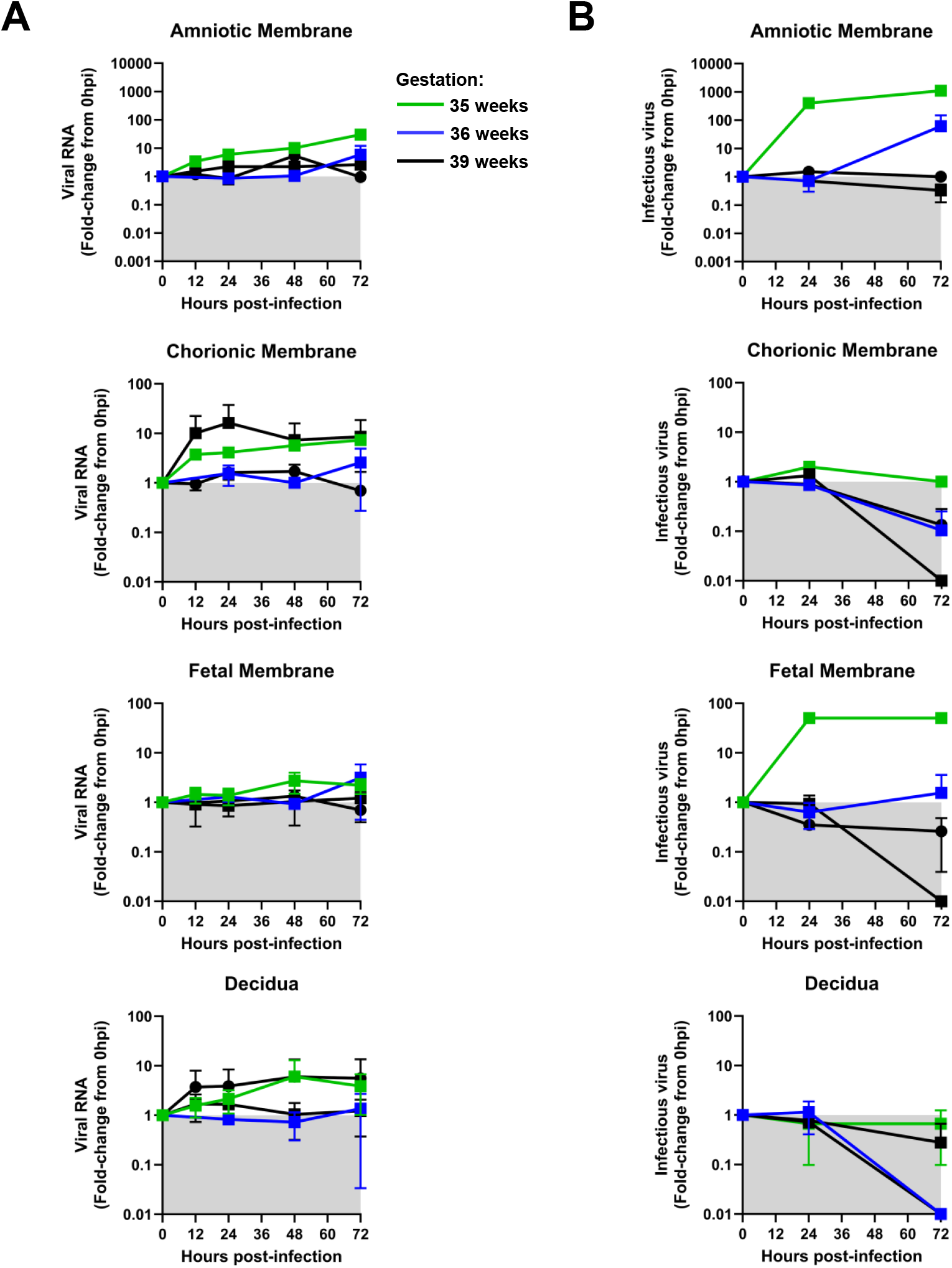
Oropouche virus infects and replicates in human fetal membrane explants. Placentas were donated at 35- (n=1), 36- (n=1), and 39-weeks (n=2) gestation. Cultures of dissected chorionic membrane, amniotic membrane, fetal membrane, and decidua were inoculated with 1 × 105 pfu of OROV (BeAn19991) for 1 hour (n=1-3 per donor). Virus was removed and washed prior to the addition of culture growth media. Culture supernatant was collected at 0, 12, 24, 48, and 72 hpi. Q-RT-PCR and viral plaque assay was performed on culture supernatant to quantify viral RNA **(A)** and infectious virus **(B)** production over time. Dashed line = LOD.

Sustained replication was not observed in chorionic membrane and decidua cultures, furthermore tissues obtained from 39-week placentas did not support OROV replication. Overall, these results indicate that OROV replicates in ex vivo placenta cultures.

## Discussion

The 2024 OROV outbreak resulted in the first suspected cases of vertical transmission in humans (3). It is not known whether this new clinical outcome is a result of increased virulence and changes in tropism, increased surveillance due to higher than average cases (32), or if vertical transmission was unrecognized during earlier outbreaks. It is feasible that it is a combination of all of these scenarios. Sequencing and phylogenetic analysis of OROV isolates collected from 2015-2024 identified the re-emergence of OROV between 2023-2024 to be associated with a novel reassortant strain (32, 35). This reassortant contained the M segment from the isolates collected in Brazil (2009-2018) and the L and S segments from isolates collected in Peru, Colombia, and Ecuador (2008-2021). *In vitro*, isolates collected from the 2024 OROV outbreak appear to be more virulent than the BeAn19991 strain in non-trophoblast cell lines (32). January 2024 the Brazilian Ministry of Health increased its epidemiological response to OROV, expanding surveillance from the Amazonas region to nationwide (36). More cases of OROV were detected after this time than from 2015-2023 (32).

Clinical evidence suggests that vertical transmission may have been previously overlooked (4). Molecular and serological analyses of historical cases of infants born with congenital malformations, but with unknown cause, identified four cases that tested positive for OROV. Each case happened during different outbreaks in Brazil prior to 2023 with one in 2014, two in 2016 and another occurring in 2018. More retrospective studies and clinical evaluations of placentas are necessary to fully appreciate the impact of OROV infection on pregnancy during and prior to the 2024 outbreak.

Our preliminary results show chorionic villous, amniotic membrane, and fetal membrane explants are susceptible to infection by a historical strain of OROV. Infection of villous explants is further supported by the high OROV titers obtained in trophoblast organoid cultures containing STB and CTB, cells that line the chorionic villous. The related bunyavirus, RVFV, also targets STB in chorionic villous explants from second and third trimester placentas (29-31). Immunofluorescence imaging of RVFV infected villi had similar virus staining patterns as OROV infected villi observed in this study and both viruses infected STB. This tropism for trophoblasts is reasonable given the surface expressed lipoprotein, Lrp1, is highly expressed on these cells and Lrp1 is a host entry factor for both OROV (37) and RVFV (38).

The lack of viral replication in the chorionic membrane in these studies may be due to local immune responses (39) dampening viral replication and may explain why less replication is found in the more complex fetal membrane, which consists of the chorionic and amniotic membranes, compared to cultures with the fetal membrane alone. Further studies identifying immune responses to OROV infection at the maternal-fetal interface are needed.

Although modest infection and replication is observed within human villous, amniotic membrane, and fetal membrane explants, the local immune response upon exposure to virus may lead to adverse pregnancy and neonatal outcomes (23, 40). As highlighted earlier, genetic changes may also contribute to increased virulence, therefore future studies should be performed comparing historical isolates to newer ones. Furthermore, five out of six samples that had positive OROV replication were from samples collected from earlier gestations (32 and 35 weeks), therefore gestation may contribute to susceptibility to infection and sustained replication. Future studies culturing additional placentas should be performed to determine whether gestation contributes to permissivity.

3D trophoblast organoids can provide a genetically tractable means to identify cellular tropism and cell-specific immune responses to infection, while explant cultures provide a means to evaluate the impact gestational age and genetic heterogeneity has on OROV infection. Explant cultures are an incomplete yet biologically relevant model due to the lack of a circulatory system, thus this model only allows us to understand the susceptibility of these structures to OROV infection in the context of its local immune response. Future studies developing an *in vivo* model are warranted to fully understand the role a complete immune response and cross-body communication plays in OROV vertical transmission.

Overall, we show that placenta explant cultures are susceptible to infection with a historical strain of OROV and syncytiotrophoblasts are among the cellular targets of the virus. Placenta explant cultures and trophoblast organoids can inform tissue and cellular tropism of OROV at the maternal-fetal interface and aid in developing countermeasures to minimize adverse pregnancy outcomes.

## Materials and methods

### Viruses

Recombinant OROV BeAn19991 (accession numbers KP052850–KP052852) was generated through reverse genetics system under the control of the T7 polymerase (41). Reverse genetics plasmids were transfected into BSR-T7 cells and culture supernatants containing infectious virus was reserved for large scale virus propagation. Virus stocks were generated by inoculating Vero E6 cells (ATCC; CL-1568) with supernatants collected from transfected BSR-T7 cells. The stock titer was determined by standard viral plaque assay (VPA). Prior to tissue or organoid inoculation, stock virus was diluted in D2 medium (DMEM, 2% (v/v) FBS, 1% L-glutamine, and 1% penicillin-streptomycin) to the desired dose. Uninfected control cultures were mock infected using D2 medium.

### Placenta procurement

Human placenta procurement was approved by the University of Pittsburgh IRB 19100322. Placentas isolated between 32-39 weeks gestation from C-section were obtained from the Steven C Caritis Magee Obstetric Maternal and Infant Biobank through an honest broker system within 30 minutes of delivery. 32-week placentas were obtained from fraternal twins (n=2). Two 39-week placentas and one 35 and 36 week placentas were also obtained for this study.

### Explant studies

Placentas were dissected within 3 hours of procurement. Dissection was conducted based on tissue type: decidua, villi, fetal membranes (chorion and/or amnion). Placenta tissue sections, approximately 5 × 5 mm sections, were placed in 24-well plates, then inoculated with 200 μL of RVFV (1.0 × 10^5^ pfu). Placenta sections were incubated for 1 hour at 37°C, 5% CO2 to allow for virus adsorption. Following the adsorption period, the inoculum was removed, tissue were washed once with 1x PBS, and then 1 mL of virus growth medium (Dulbecco’s modified Eagle medium (DMEM), 2% (vol/vol) fetal bovine serum (FBS), 1% L-glutamine, and 1% penicillin– streptomycin) was added to all samples. The tissues were cultured for 72 hours and 100 μL of culture supernatant was pooled from two wells from each tissue daily at 0, 12, 24, 48, and 72 hpi. For each human placenta donor, 1-3 pooled samples were analyzed from each time point.

### 3D trophoblast organoid studies

Appropriately polarized 3D trophoblast organoids were generated as previously described (33). Briefly CT-27 trophoblast organoids were grown on functionalized polystyrene beads for six days, as previously described^1^. Exceptions to this protocol are as follows: CT27 seeding was performed at 40,000 cells/mL, and organoids were kept in cTOM media (1x ITS-X, 1% KSR, 0.15% BSA, 5 μL/mL Penicillin-Streptomycin, 200 μM L-ascorbic acid, 40 μM 2-mercaptoethanol, 2 μM CHIR99021, 5 μM A83-01, 2.5 μM Y27632, 25 ng/mL hEFG, and 37.5 uM VPA) for the first 72 hours before transitioning to TOM media for 3 additional days to induce syncytialization.

Organoids were placed in a microcentrifuge tube and centrifuged for 3 minutes at 2,000 rpm to remove culture supernatant. 100 μL of stock OROV BeAn19991 (1.0 × 10^5^ pfu) was added to each tube containing organoids, then incubated at 37°C, 5% CO2 for 1 hour to allow for virus adsorption. Following the adsorption period, the microcentrifuge tube was centrifuged to remove the virus inoculum and washed three times with 1x PBS. Then 1mL of TOM was added to all samples then returned to the corresponding wells of a 6-well plate. The plate was cultured for 48 hours and 200 μL was collected from each well at 0, 12, 24, 36, and 48 hpi. Four (n=4) wells were inoculated with OROV, and two (n=2) wells were mock inoculated with D2 medium.

### RNA isolation and q-RT-PCR

The culture supernatant (50 μL) was inactivated in 900 μL of TRIzol reagent (Invitrogen). RNA isolation and q-RT-PCR were performed as describe (42). OROV BeAn19991-specific primers and probes were used for q-RT-PCR: forward primer – 5’ TACCCAGATGCGATCACCAA 3’, reverse primer – 5’ TTGCGTCACATCATTCCAA 3’, probe 5’ 6-FAM-TGCCTTTGGCTGAG GTAAAGGGCTC – BHQ_1 3’. Semi-quantitation of virus was determined by comparing cycle threshold (CT) values from unknown samples to CT values from the in-house developed OROV BeAn19991 RNA standards based on PFU equivalents per mL

### Immunohistochemistry, immunofluorescence, and microscopy

For immunohistochemistry (IHC), explant tissues were fixed in 4% PFA for 24 hours then paraffin embedded by standard protocols. IHC was performed as explained in McMillen et al. (30) using anti-OROV nucleoprotein IgG (1:100 dilution; custom made by Genscript). Primary delete and no infection controls were performed to identify non-specific binding of the secondary detection antibody or all reagents, respectively. Red immunopositivity observed above the no infection control was deemed as positive for OROV antigen. Colorimetric ISH slides were imaged using an Olympus CX41 microscope with the Levenhuk M300 base attachment. Denoising and contrasting were performed using the Adobe Illustrator.

For immunofluorescence, fixed organoids and tissue were permeabilized with 0.25% Triton X-100 for 15 minutes at room temperature and placed in 1% bovine serum albumin for 1 hour.

Anti-OROV antibodies were incubated for one hour at 1:500 dilution followed by goat-anti rat antibody conjugated to Alexa-488 for 30 minutes at room temperature. J2 conjugated antibody was used at a 1:1000 dilution. Phalloidin 488 or 568 was incubated for 30 minutes. Organoids were then mounted using Vectashield mounting medium and imaged using a Nikon confocal microscope at the Magee Womens Research Institute core facility.

## Acknowledgements

All data needed to evaluate the conclusions in the paper are present in the paper and/or the Supplementary Materials. Additional data related to this paper may be requested from the authors. The authors declare that they have no competing interests. All work was supported by the National Institutes of Health (K01 AI165965 to C.M.M, R01 AI150792 and R01 AI150792S1 to A.L.H.). The funders had no role in study design, data collection and analysis, decision to publish, or preparation of the manuscript. Conceptualization: Cynthia M. McMillen; Data curation: Cynthia M. McMillen, Christina Megli, Jackson J. McGaughey. Formal analysis: Cynthia M. McMillen, Christina Megli, Amy L. Hartman; Funding acquisition: Cynthia M. McMillen, Amy L. Hartman; Investigation: Cynthia M. McMillen, Christina Megli, Rebecca K Zack, Jackson J. McGaughey, Ryan M. Hoehl, Amy L. Hartman; Methodology: Cynthia M. McMillen, Christina Megli, Jackson J. McGaughey, Ryan M. Hoehl, Taylor Snisky; Project administration: Cynthia M. McMillen, Amy L. Hartman; Supervision: Cynthia M. McMillen; Validation: Cynthia M. McMillen; Visualization: Cynthia M. McMillen, Christina Megli; Writing – original draft: Cynthia M. McMillen, Christina Megli.

## Figure Legend

**Supplemental Figure 1:**
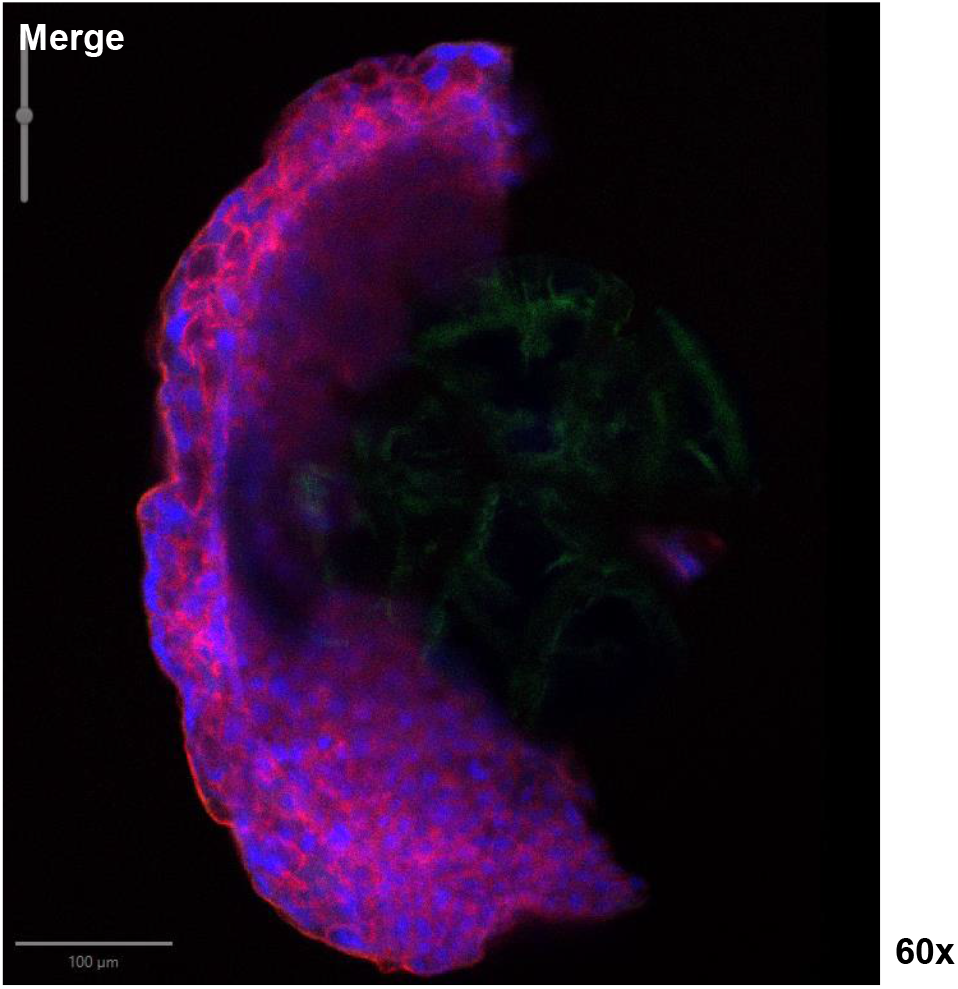
Immunofluorescence images of uninfected organoid control. Blue = DAPI, red = phalloidin, green = OROV nucleoprotein (N). Image was taken at 60x magnification.

